# *Le Petit Prince*: A multilingual fMRI corpus using ecological stimuli

**DOI:** 10.1101/2021.10.02.462875

**Authors:** Jixing Li, Shohini Bhattasali, Shulin Zhang, Berta Franzluebbers, Wen-Ming Luh, R. Nathan Spreng, Jonathan R. Brennan, Yiming Yang, Christophe Pallier, John Hale

## Abstract

Neuroimaging using more ecologically valid stimuli such as audiobooks has advanced our understanding of natural language comprehension in the brain. However, prior naturalistic stimuli have typically been restricted to a single language, which limited generalizability beyond small typological domains. Here we present the *Le Petit Prince* fMRI Corpus (LPPC–fMRI), a multilingual resource for research in the cognitive neuroscience of speech and language during naturalistic listening (Open-Neuro: ds003643). 49 English speakers, 35 Chinese speakers and 28 French speakers listened to the same audiobook *The Little Prince* in their native language while multi-echo functional magnetic resonance imaging was acquired. We also provide time-aligned speech annotation and word-by-word predictors obtained using natural language processing tools. The resulting timeseries data are shown to be of high quality with good temporal signal-to-noise ratio and high inter-subject correlation. Data-driven functional analyses provide further evidence of data quality. This annotated, multilingual fMRI dataset facilitates future re-analysis that addresses cross-linguistic commonalities and differences in the neural substrate of language processing on multiple perceptual and linguistic levels.

## Background & Summary

In the cognitive neuroscience of language, there is a growing consensus that using more ecologically valid stimuli such as audiobooks might extend our understanding of language processing in the brain^1–3^. Compared to traditional factorial designs with a large number of repetitive trials, naturalistic paradigms use stories and dialogues with a rich context and produce results that are generalizable to everyday language use^3,4^. However, prior naturalistic studies have typically been restricted to a single language, which limited neurobiological frameworks for language processing to small typological domains. Here we present *Le Petit Prince* fMRI Corpus (LPPC-fMRI), a multilingual fMRI dataset where English, Chinese and French speakers listened to the same audiobook *Le Petit Prince (The Little Prince)* in their native language (see Figure 1 for a Schematic overview of the LPPC-fMRI data collection, preprocessing, technical validation and annotation procedures). Our parallel corpus facilitates future research on cross-linguistic commonalities and differences in the neural processes for language comprehension.

**Figure 1.**
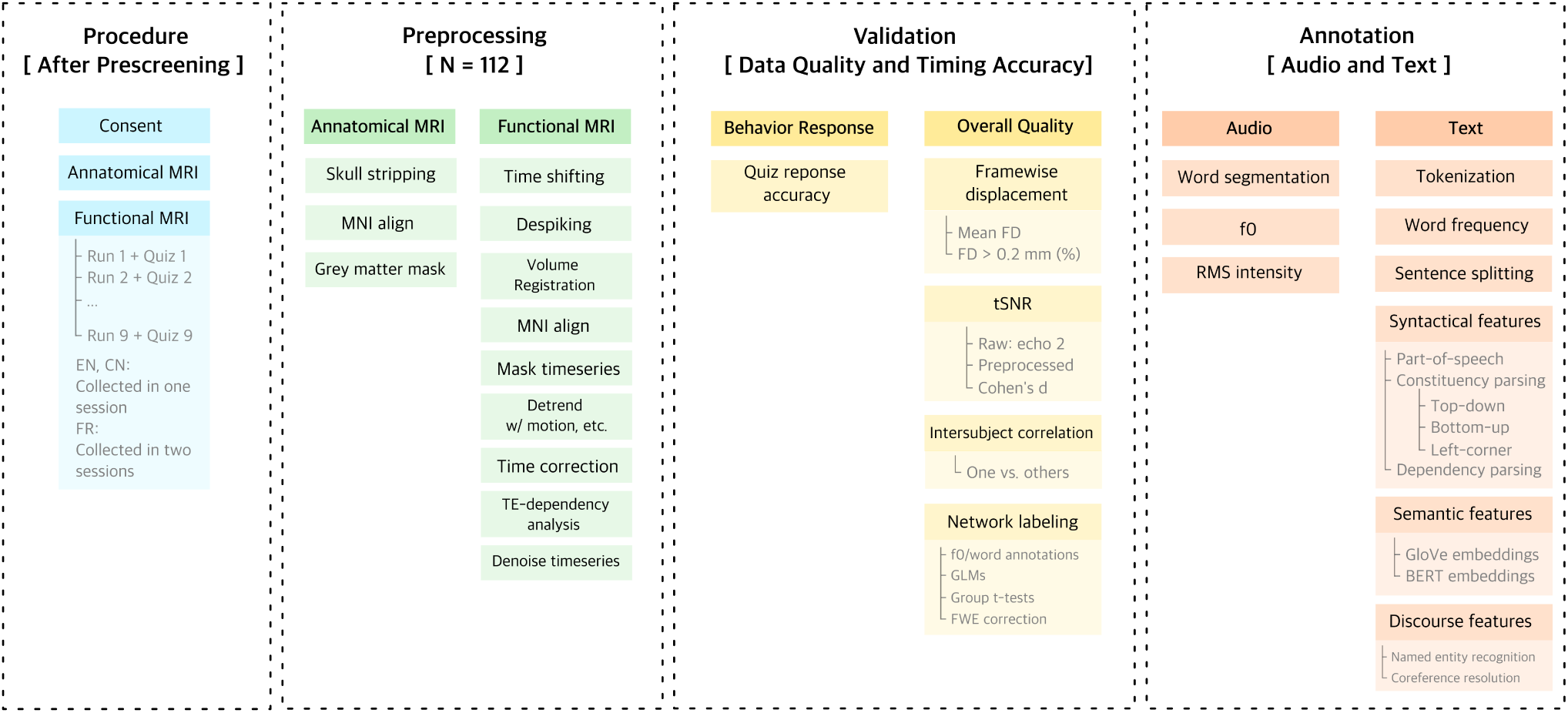
Schematic overview of the LPPC-fMRI data collection procedures, preprocessing, technical validation and annotation. During data collection (blue), anatomical MRI was first acquired, followed by functional MRI while participants listened to 9 sections of the audiobook. After preprocessing the data (green), behavioral and overall data quality were examined (yellow). Audio and text annotations were extracted using NLP tools.

In naturalistic designs such as story listening, linguistic processes on multiple levels (e.g., word, phrase, sentence, discourse) unfold naturally at different timescales. Such a rich contextual setting extends the range of linguistic phenomena that can be examined in parallel, and allows for testing assumptions on the neural mechanisms of language processing. For example, whether different linguistic levels coincide with different frequencies of oscillatory activity in the brain^5,6^, and whether these levels correspond to a hierarchically organized predictive coding architecture^7^. In addition, naturalistic approaches to neurolinguistics are in synergy with natural language processing (NLP), where using ecologically valid language corpora for training models has been common practice for the past quarter-century. Accordingly, NLP models can be leveraged to understand linguistic processes at an algorithmic level by comparing model predictions against brain data during naturalistic comprehension. For example, syntactic structure-building as predicted by the bottom-up or left-corner parsing strategies^8–10^ and recurrent neural network grammars (RNNG)^11^ has been shown to fit well with left temporal activity. Recent neural network architectures such as bidirectional LSTMs^12^ and Transformers^13^ have also been shown to correlate with neural responses during naturalistic comprehension, suggesting construction-specific variations in the understanding of linguistic expressions.

While naturalistic designs opened up a host of new research questions that are not possible to study under tightly controlled experimental designs, the majority of prior naturalistic studies have been restricted to a single language. This limited our understanding of the neural processes of language comprehension to small typological domains. To complement monolingual datasets such as the Narrative Brain Dataset (NBD)^14^, the Alice Dataset^15^ and the Mother of Unification Studies^16^, we collected a multilingual fMRI dataset consisted of Antoine de Saint-Exupéry’s *The Little Prince* in English, Chinese and French. A total of 112 subjects (49 English speakers, 35 Chinese speakers and 28 French speakers) listened to the whole audiobook for about 100 minutes in the scanner (see Table 1 and Table 2 for the demographics of the participants, data collection procedures, and stimuli information for the English, Chinese, and French datasets). This stimulus is considerably longer than other datasets (i.e., 6 minutes on average for the NBD dataset and 12 minutes for the Alice dataset), allowing for testing linguistic phenomena that may not be sufficiently attested in smaller samples. This dataset includes time-aligned speech segmentation, prosodic information and word-by-word predictors obtained using natural language processing tools, ranging from lexical semantics to syntax to discourse information (see Figure 2 for the annotations available for an example sentence from the English audiobook). The neuroimaging data, as well as the annotations and information about the experimental procedure are shared in a standardized format on OpenNeuro^17^.

**Table 1.** Demographics of the participants, data collection procedures, and stimuli information for the English, Chinese, and French datasets.

| Language | Participants |  |  | Data Collection |  | Stimuli |  |  |
| --- | --- | --- | --- | --- | --- | --- | --- | --- |
|  | Number | Mean Age | Female | Location | Material | Length (s) | N Words | N Sentences |
| English | 49 | 21.3 | 30 | Cornell University, United States | The little prince EN audiobook | 5632 | 15376 | 1499 |
| Chinese | 35 | 19.9 | 15 | Jiangsu Normal University, China | The little prince CN audiobook | 5954 | 16009 | 1577 |
| French | 28 | 24.4 | 15 | NeuroSpin, France | The little prince FR audiobook | 5828 | 15391 | 1480 |

**Table 2.**
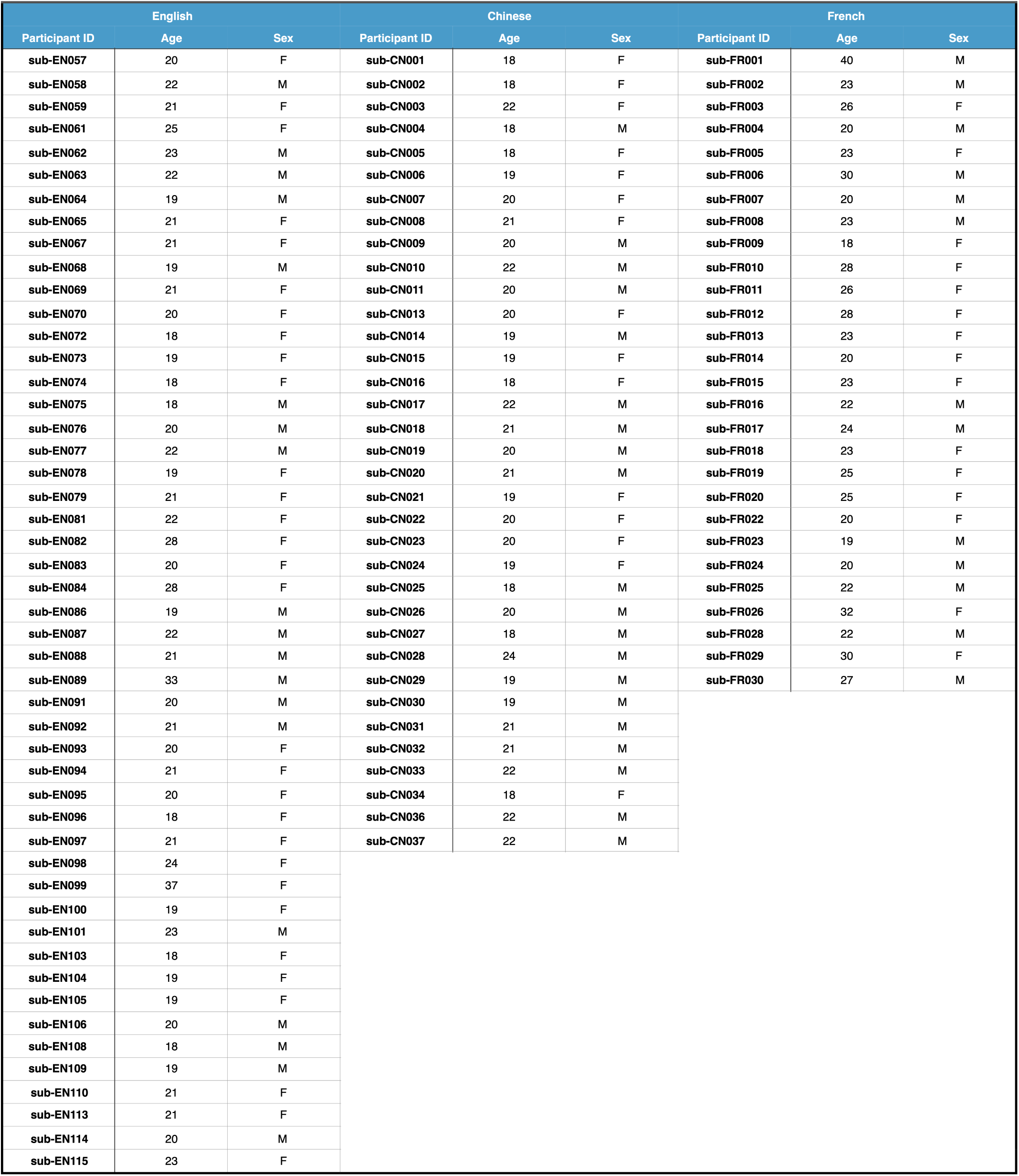
List of subjects in the data collection with basic demographic information.

**Figure 2.**
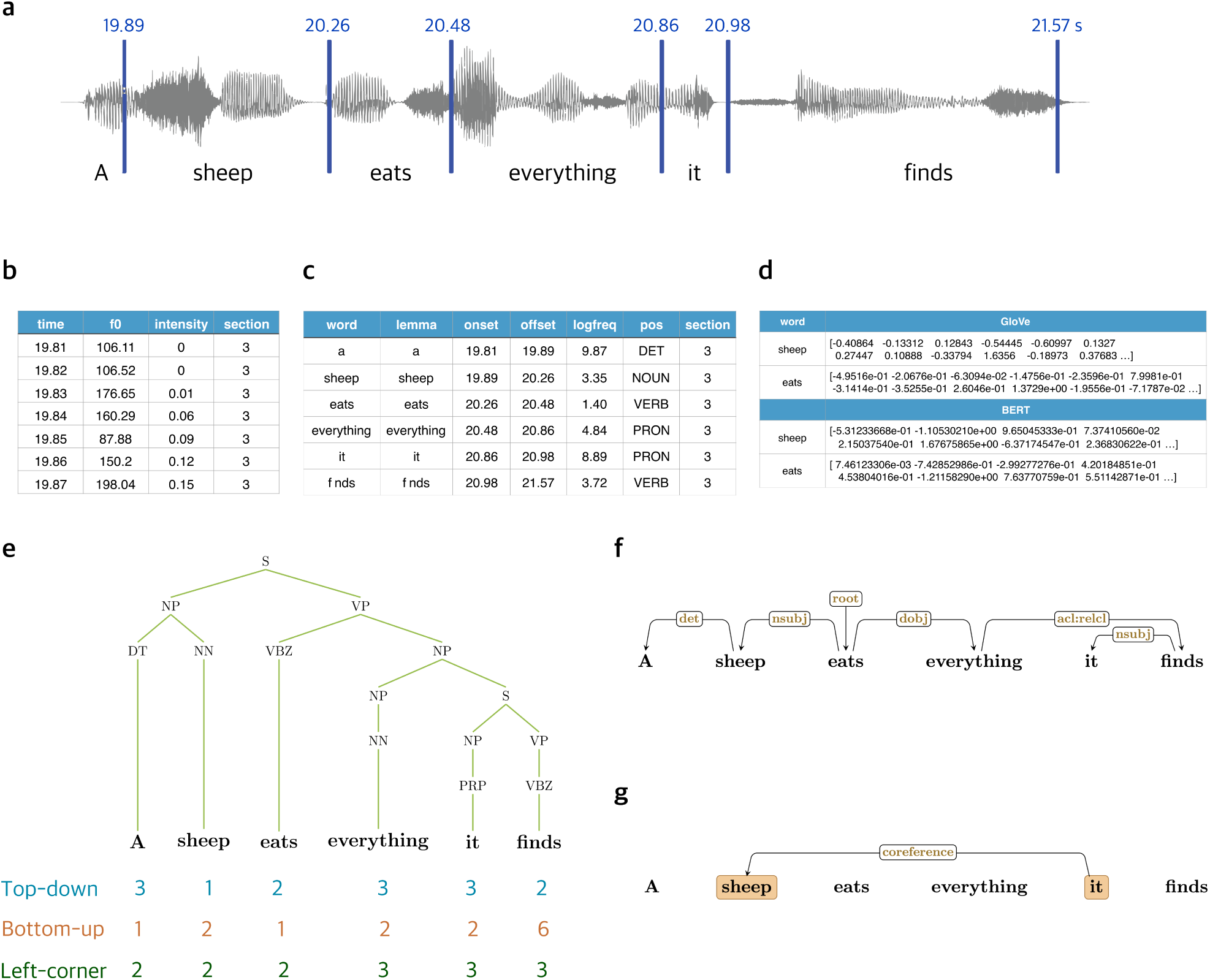
Annotation information for the stimuli. **a**. Word boundaries in the audio files, included in files: lpp<EN/CN/FR>_section[1-9].TextGrid. **b**. f0 and RMS intensity for every 10 ms of the audios, included in files: lpp<EN/CN/FR>_prosody.csv **c**. Tokenization, lemmatization, log-tranformed word frequency and POS tagging, included in files: lpp<EN/CN/FR>_word_information.csv. **d**. GloVe and BERT embeddings for every word in the audiobooks, included in files: lpp<EN/CN/FR>_word_embeddings_GloVe.csv and lpp<EN/CN/FR>_word_embeddings_BERT.csv **e**. Parsed syntactic trees based on constituency grammar with node counts using top-down, bottom-up, and left-corner parsing strategies^31^, included in files: lpp<EN/CN/FR>_trees.csv **f**. Dependency relations for each words in each sentence, included in files: lpp<EN/CN/FR>_dependency.csv. **g**. Named entity recognition and coreference relations for the English and Chinese texts, included in files: lpp<EN/CN>_coreference.csv.

**Figure 3.**
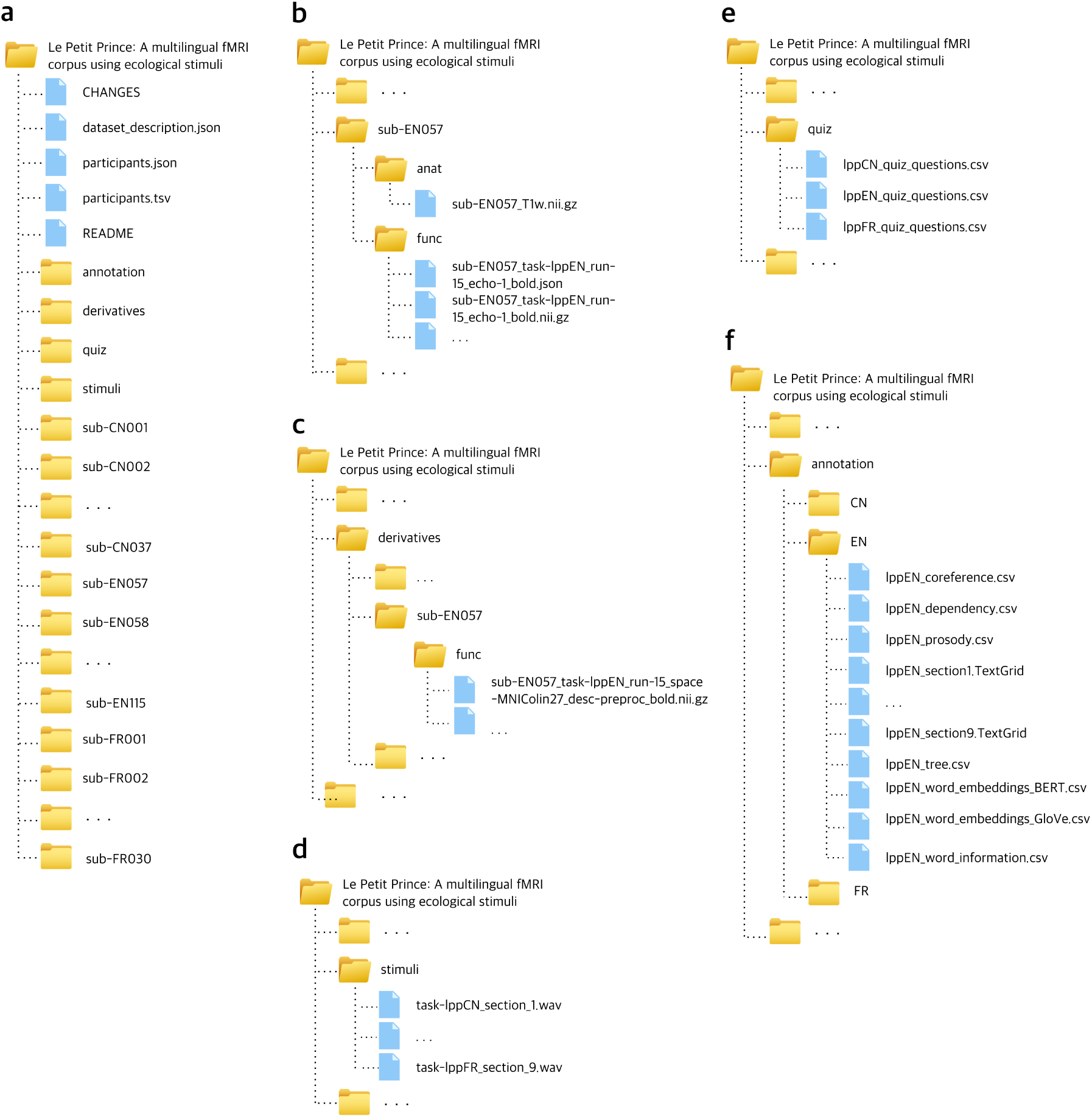
Organization of the data collection. **a**. General overview of directory structure. **b**. Content of subject-specific anatomical and raw data directories. **c**. Content of subject-specific preprocessed data directories. **d**. Content of the stimuli directory. **e**. Content of the quiz directory. **f**. Content of the language-specific annotation directory.

The LPPC-fMRI facilitates cross-linguistic generalization and helps overcome current statistical and typological limitations in the neurobiology of language. We stress the importance of considering multiple languages when building and testing neurobiological models of language processing, assuming that the neural substrates and processes of language are shared among speakers of all languages. As shown in previous work examining coreference resolution using the English and Chinese subset of this corpus, the computational model that best explains the neural signature for pronoun processing is generalizable for both English and Chinese^18^. These data can be reused to address different research questions with a variety of analytical methods. Future work envisions an expanded LPPC, one that incorporates data from additional neuroimaging modalities, such as electrocorticography (EEG) and magnetoencephalography (MEG). For instance, LPPC-EEG dataset aspires to 26 languages^4^. Our vision is for the LPPC to become an open infrastructure to which researchers from various communities can contribute by adding further modalities, languages and annotations.

## Methods

### Participants

A total of 112 subjects listened to the whole audiobook for about 100 minutes in the scanner. Table 1 and Table 2 show the summary of the data collection procedure, the stimuli and participants information for the three datasets.

English participants were 49 young adults (30 females, mean age=21.3, SD=3.6) with no history of psychiatric, neurological or other medical illness that might compromise cognitive functions. They self-identified as native English speakers, and strictly qualified as right-handed on the Edinburgh handedness inventory^19^. All participants were paid, and gave written informed consent prior to participation, in accordance with the IRB guidelines of Cornell University.

Chinese participants were 35 healthy, right-handed young adults (15 females, mean age=19.3, SD=1.6). They self-identified as native Chinese speakers, and had no history of psychiatric, neurological, or other medical illness that could compromise cognitive functions. All participants were paid, and gave written informed consent prior to participation, in accordance with the IRB guidelines of Jiangsu Normal University.

French participants were 28 healthy, right-handed adults (15 females, mean age=24.4, SD=4.6). They self-identified as native French speakers and had no history of psychiatric, neurological, or other medical illness that could compromise cognitive functions. All participants gave written informed consent prior to participation, in accordance with the Regional Committee for the Protection of Persons involved in Biomedical Research.

### Procedures

After giving their informed consent, participants were familiarized with the MRI facility and assumed a supine position on the scanner. They were instructed to not move as best as they could throughout scanning as movement would make the scans unusable. Next, participants were put in the head-coil with pillows under and on the sides of their head and under the knees for comfort and to reduce movement over the scanning session. Participants were given a bulb in their right hand and told to squeeze if something was wrong or they needed a break during scanning. Once in place, participants chose an optimal stimulus volume by determining a level that was loud but comfortable. Auditory stimuli were delivered through MRI-safe, high-fidelity headphones inside the head coil (English: Confon HP-VS01, MR Confon, Magdeburg, Germany; Chinese: Ear Bud Headset, Resonance Technology, Inc, California, USA; French: Magnacoil TIM headset, Siemens, Germany). The headphones were secured against the plastic frame of the coil using foam blocks.

The English and Chinese participants went through one scanning session, which was divided into 9 runs, and each lasted for about 10 minutes. Participants listened passively to 1 section of the audiobook in each run and completed 4 quiz questions after each run (36 questions in total). These questions were used to confirm their comprehension and were viewed by the participants via a mirror attached to the head coil and they answered through a button box. During scanning, participants were monitored by a camera over their left eye. If they appeared drowsy or seemed to move too much during the movie, the operator of the scanner gave them a warning over the intercom by producing a beep or speaking to them. During breaks between the runs, participants were told that they could relax but not move. Finally, participants were paid and sent home. The entire session lasted for around 2.5 hours. In French, due to a legal limitation, participants could not stay for longer than 1.5 hours inside the scanner; therefore, the acquisition was split into two sessions separated by a period of 1 to 2 hours out of the scanner.

### Stimuli

The English *The Little Prince* audiobook is 94 minutes long, translated by David Wilkinson and read by Karen Savage. The Chinese audiobook^20^ is 99 minutes long, read by a professional female Chinese broadcaster hired by the experimenter. The French audiobook is 97 minutes long, read by Nadine Eckert-Boulet and published by the now-defunct Omilia Languages Ltd. The original French text is copyrighted by Gallimard 1946.

### Acquisition

English and Chinese MRI images were acquired with a 3T MRI GE Discovery MR750 scanner with a 32-channel head coil. French MRI images were acquired with a 3T Siemens Magnetom Prisma Fit 230 scanner. Anatomical scans were acquired using a T1-weighted volumetric Magnetization Prepared Rapid Gradient-Echo (MP-RAGE) pulse sequence. Functional scans were acquired using a multi-echo planar imaging (ME-EPI) sequence with online reconstruction (TR=2000 ms; English and Chinese: TEs=12.8, 27.5, 43 ms; French: TEs=10, 25, 38 ms; FA=77°; matrix size=72 × 72; FOV=240.0 mm × 240.0 mm; 2 × image acceleration; English and Chinese: 33 axial slices; French: 34 axial slices; voxel size=3.75 × 3.75 × 3.8 mm).

### Preprocessing

MRI data files were converted from DICOM to NIfTI format and preprocessed using AFNI version 16^21^.

#### Anatomical

The anatomical/structural MRI scans were deskulled using *3dSkullStrip*. The resulting anatomical images were nonlinearly aligned to the Montreal Neurological Institute (MNI) N27 template brain. Resulting anatomical images were used to create grey matter masks.

#### Functional

The first 4 volumes in each run were excluded from analyses to allow for T1-equilibration effects. The fMRI time-series were then corrected for slice-timing differences (*3dTshift*) and despiked (*3dDespike*). Next, volume registration was done by aligning each timepoint to the mean functional image of the centre timeseries (*3dvolreg*). Then the volume-registered and anatomically-aligned functional data were nonlinearly aligned to the MNI template brain. Multi-echo independent components analysis (ME-ICA)^22^ were used to denoise data for motion, physiology and scanner artifacts. Images were then resampled at 2 mm cubic voxels (*3dresample*).

### Annotations

Apart from the fMRI timeseries data, we also provide audio and text annotations ranging from time-aligned speech segmentation and prosodic information to word-by-word predictors obtained using natural language processing tools, including lexical semantics, syntax and discourse-level information. See Figure 2 for a summary of our annotations. These annotations are available on OpenNeuro too (see the Data records section).

#### Speech segmentation

Word boundaries in the audio were identified and aligned to the transcripts using Forced Alignment and Vowel Extraction (FAVE)^23^ and were manually checked by two native speakers of the three languages.

#### Prosodic information

Root mean square intensity and the fundamental frequency (f0) for every 10 ms of each audio section of the three languages were extracted using the Voicebox toolbox^24^.

#### Word frequency

Log-transformed unigram frequency of each word in *The Little Prince* in English, Chinese and French was estimated using Google Books Ngram Viewer, Version 20120701^25^.

#### Word embeddings

Static GloVe embeddings^26^ and contextualized BERT embeddings for each word (given its sentential context) in the *The Little Prince* in the three languages were extracted using the SpaCy package^27^. Words that are divided into subwords by BERT used the average embedding of the subwords.

#### Part-of-speech tagging

Part-of-speech (POS) tagging for each word in the book in the three languages was extracted using the Stanford parser for English^28^, Chinese^29^ and French^30^.

#### Constituency parsing

Syntactic tree structures of each sentence in the audiobooks was parsed using the Stanford parser for English^28^, Chinese^29^ and French^30^.

#### Parser actions

Syntactic node counts for each word in the audiobooks based on bottom-up, top-down and left-corner parsing strategies^31^ as applied to the Stanford-derived constituency trees described above. These word-by-word counts are the number of parser actions that would be taken (on a given strategy) before moving on to the next word in the sentence. They were calculated using custom tree-walking software.

#### Dependency parsing

Dependency relations of words in each sentence of the audiobooks were parsed using the Stanford dependency parser for English^32^, Chinese^33^ and French^30^.

#### Coreference resolution

Antecedents for each third person pronoun in the English and Chinese audiobooks were manually annotated using the annotation tool brat^34^.

### Data records

Information and anatomical data that could be used to identify participants has been removed from all records. Resulting files are available from the OpenNeuro platform at https://openneuro.org/datasets/ds003643. A README file there provides a description of the available content. The code/scripts used for this manuscript are available on GitHub (https://github.com/jixing-li/lpp_data).

### Participant responses

**Location** participants.json, participants.tsv

**File format** tab-separated value

Participants’ sex, age and responses to quiz questions in tab-separated value (tsv) files. Data is structured as one line per participant.

### Audio files

**Location** stimuli/task-lpp<EN/CN/FR>_section_[1-9].wav

**File format** wav

The English, Chinese and French audiobooks divided into nine sections.

### Anatomical MRI

**Location** sub-<EN/CN/FR><ID>/anat/sub-<EN/CN/FR><ID>_T1w.nii.gz

**File format** NIfTI, gzip-compressed

The defaced raw high-resolution anatomical image.

### Functional MRI

**Location** sub-<EN/CN/FR><ID>/func/sub-<EN/CN/FR><ID>_task-lpp<EN/CN/FR>_run-0[1-9]_echo-[1-3]_bold.nii.gz

**File format** NIfTI, gzip-compressed

**Sequence protocol** sub-<EN/CN/FR><ID>/func/sub-<EN/CN/FR><ID>_task-lpp<EN/CN/FR>_run-0[1-9]_echo-[1-3]_bold.json

The mutli-echo fMRI data are available as individual timeseries files, stored as sub-<EN/CN/FR><ID>/func/sub-<EN/CN/FR><ID>_task-lpp<EN/CN/FR>_run-0[1-9]_echo-[1-3]_bold.nii.gz. The MEI-CA preprocessed timeseries are also available as derivatives/sub<EN/CN/FR><ID>/func/sub-<EN/CN/FR><ID>_task-lpp<EN/CN/FR>_run-0[1-9]_space-MNIColin27_desc-preproc_bold.nii.gz.

### Annotations

**Location** annotation/<EN/CN/FR>/lpp<EN/CN/FR>_section[1-9].TextGrid,

annotation/<EN/CN/FR>/lpp<EN/CN/FR>_prosody.csv,

annotation/<EN/CN/FR>/lpp<EN/CN/FR>_word_information.csv,

annotation/<EN/CN/FR>/lpp<EN/CN/FR>_word_embeddings_GloVe.csv,

annotation/<EN/CN/FR>/lpp<EN/CN/FR>_word_embeddings_BERT.csv

annotation/<EN/CN/FR>/lpp<EN/CN/FR>_tree.csv,

annotation/<EN/CN/FR>/lpp<EN/CN/FR>_dependency.csv,

annotation/<CN/EN>/lpp<CN/EN>_coreference.csv

**File format** comma-separated value

Speech and linguistic annotations for the audio and text of the three languages.

### Quiz questions

**Location** quiz/lpp<EN/CN/FR>_quiz_questions.csv

**File format** comma-separated value

The 36 comprehension quiz questions used in the English, Chinese and French experiments.

### Technical Validation

Accuracy of participants’ responses to the quizzes after each section was calculated to ensure adequate comprehension. To assess fMRI scan quality, we calculated framewise displacement (FD), temporal signal-to-noise ratio (tSNR) and inter-subject correlation (ISC). We also did two whole-brain functional analyses using pitch (f0) and word annotations. These serve to show data quality similar to past work and provide evidence for timing accuracy between fMRI timeseries for participants.

### Behavioral results

Participants answered four four-choice comprehension questions after each section (36 questions in total). An example question is shown below. Participants performed well with a mean accuracy of 89.5% (SD=3.8) and 86.4% (SD=2.7) for English and Chinese participants, respectively. French participants responses were noted on paper by the experimenters during recording and were unfortunately unable to locate now. But the experimenters did not notice any French participant with an abnormally low accuracy (<75%) for the quiz questions.

Why was the little prince difficult to talk toã

a. He spoke a foreign language.
b. He was mute.
c. He didn’t ask enough questions.
d. He didn’t answer questions directly.

Key: (d)

### Framewise displacement

Framewise displacement is a measure of the frame-to-frame movement, assessed in millimetres. The six motion parameters (3 translation parameters and 3 rotation parameters) generated by MEI-CA.py were used to calculate FD, defined as the sum of the absolute temporal derivatives of the six motion parameters, following conversion of rotational parameters to distances by computing the arc length displacement on the surface of a sphere with radius 50 mm^35,36^:

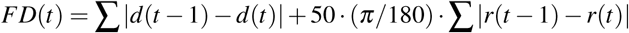

where d denotes translation distances *x, y, z*, and r denotes rotation angles *α, β, γ*. For each participant, a single (scalar) estimate of overall motion, the mean FD, can be calculated by averaging the FD time series.

For the English data, the average FD was 0.11 mm (SD=0.05); for the Chinese data, the average FD was 0.08 mm (SD=0.05), and for the French data, the average FD was 0.10 mm (SD=0.02). FD values greater than 0.20 mm are conventionally considered high motion^36^, we therefore also calculated the percentage of frames for each subject where FD exceeded 0.20 mm. The average percentage of frames where FD was greater than 0.20 mm was 9.3% (SD=10.6%), 5.0% (SD=8.2%) and 4.6% (SD=5.0%) for the English, Chinese and French data, respectively (see Table 3).

**Table 3.** Summary of framewise displacement information for the English, Chinese and French data.

|  | FD (mm) |  | FD > 0.2 mm (%) |  |
| --- | --- | --- | --- | --- |
|  | Mean | SD | Mean | SD |
| English | 0.11 | 0.05 | 9.3 | 10.6 |
| Chinese | 0.08 | 0.05 | 5.0 | 8.2 |
| French | 0.10 | 0.02 | 4.6 | 5.0 |

### Temporal signal-to-noise ratio

tSNR is a measure of signal strength at the voxel level, defined as the mean signal intensity of a voxel across the timeseries divided by its standard deviation. We calculated tSNR both before preprocessing using the middle echo image which most closely approximates standard single echo collection, and after the optimal combination of the echo images with MEI-CA denoising. We compared the tSNR values before and after extensive preprocessing using Cohen’s d:

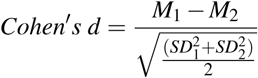

where M and SD are the mean and standard deviation of the tSNR in a voxel for the more (subscript one) minus the less preprocessed timeseries (subscript two). We applied a grey matter mask with most white matter and ventricle voxels removed. The tSNR values showed a clear increase after MEI-CA denoising across the three language groups, suggesting clearer signal compared to standard single echo acquisition (see Figure 4).

**Figure 4.**
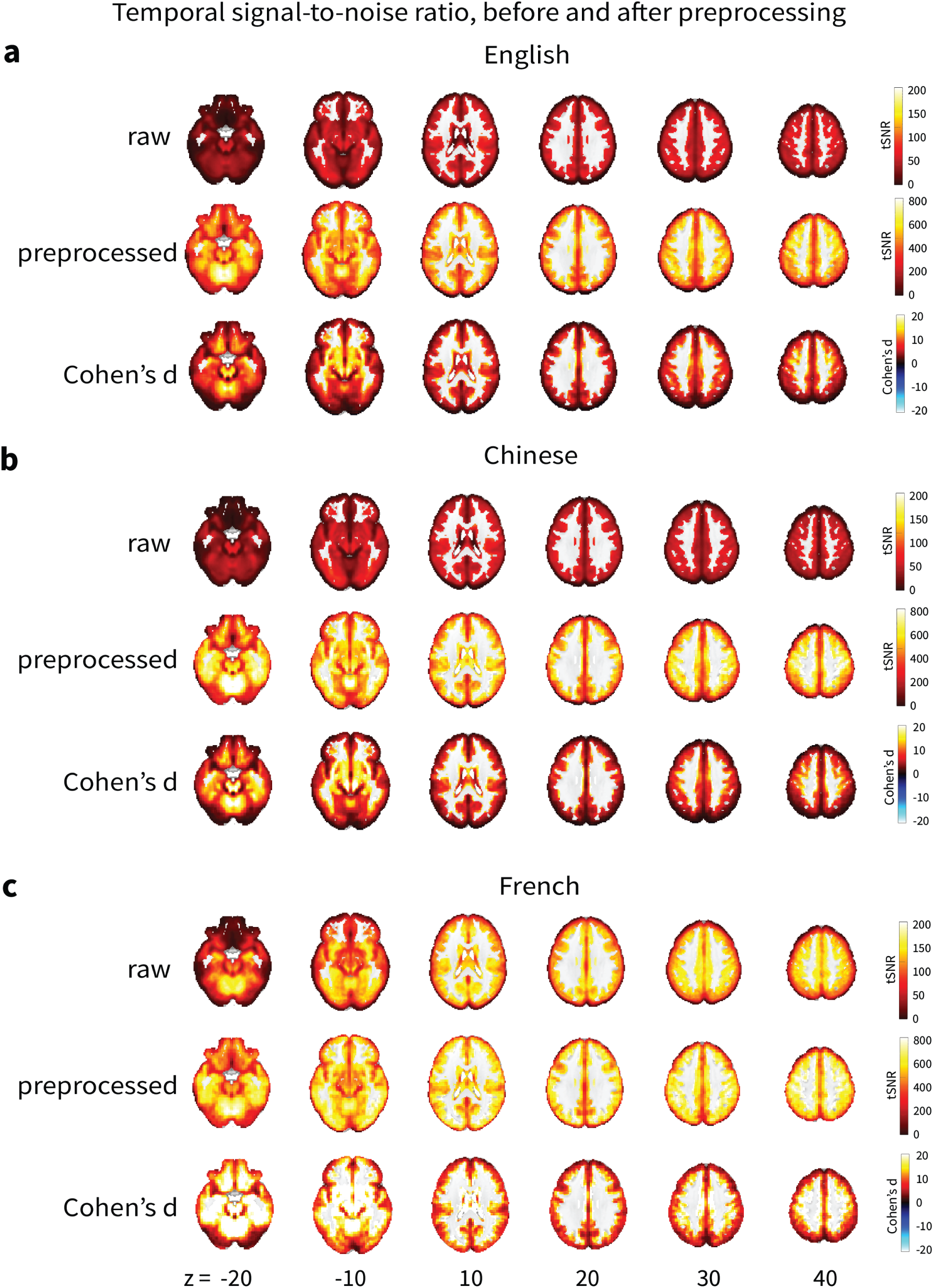
Voxel-wise temporal signal-to-noise ratio analysis before and after preprocessing. Cohen’s d effect sizes showed increase in tSNR after preprocessing.

### Inter-subject correlation

To estimate what proportion of the brain signal in response to the audiobook was consistent across subjects, we computed the inter-subject correlation (ISC) for each voxel’s timeseries across subjects in each language group. Each subjects data in a voxel was correlated to the average timeseries of the other subjects in the same voxel. This generated a map that quantifies the similarity of an individual subjects response with the group response. The procedure was repeated for all subjects, and a median ISC map was computed at the group level. The ISC results showed largest correlation in brain responses across subjects in the temporal regions, the brain regions implicated for speech and language processing (see Figure 5).

**Figure 5.**
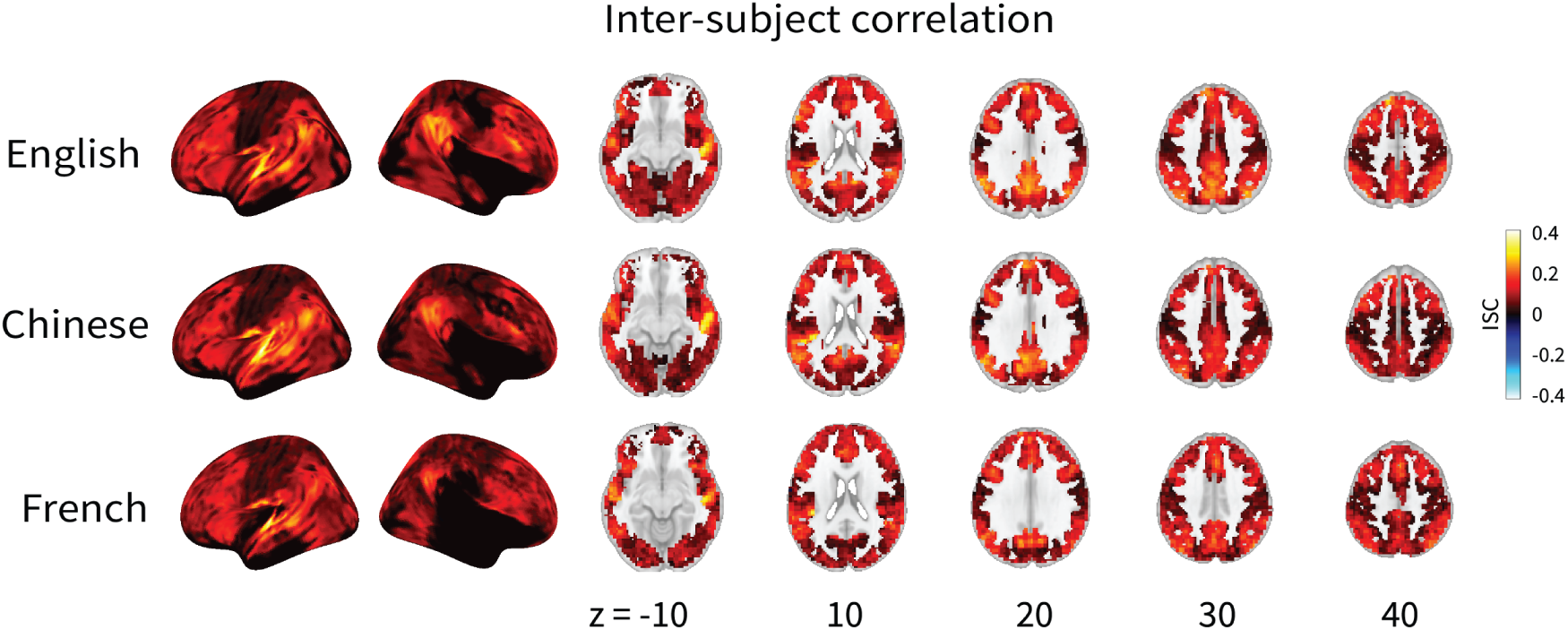
Results of inter-subject correlation (ISC) demonstrating data quality and timing synchrony between participants. As expected, the temporal regions showed the largest correlation in brain responses across subjects

### Network labeling

Besides demonstrating data and timing quality, here we also illustrate the general linear model (GLM) methods to derive the prosody and word regions using our pitch and word annotations. In particular, we calculated the *f0* for every 10 ms of the audio in each language and marked 1 at the offset of each word in the audio (*wordrate*). We then convolved the *f0* and *wordrate* annotations with a canonical hemodynamic response function and regressed them against the preprocessed fMRI timecourses using GLMs. At the group level, the contrast images for the *f0* and *wordrate* regressors were examined by a one-sample *t*-test. An 8 mm full-width at half-maximum (FWHM) Gaussian smoothing kernel was applied on the contrast images from the first-level analysis to counteract inter-subject anatomical variation. Statistical significance was held at *p <* .05 FWE with a cluster size greater than 50. Figure 6 illustrates the GLM methods to localize the pitch and word regions.

**Figure 6.**
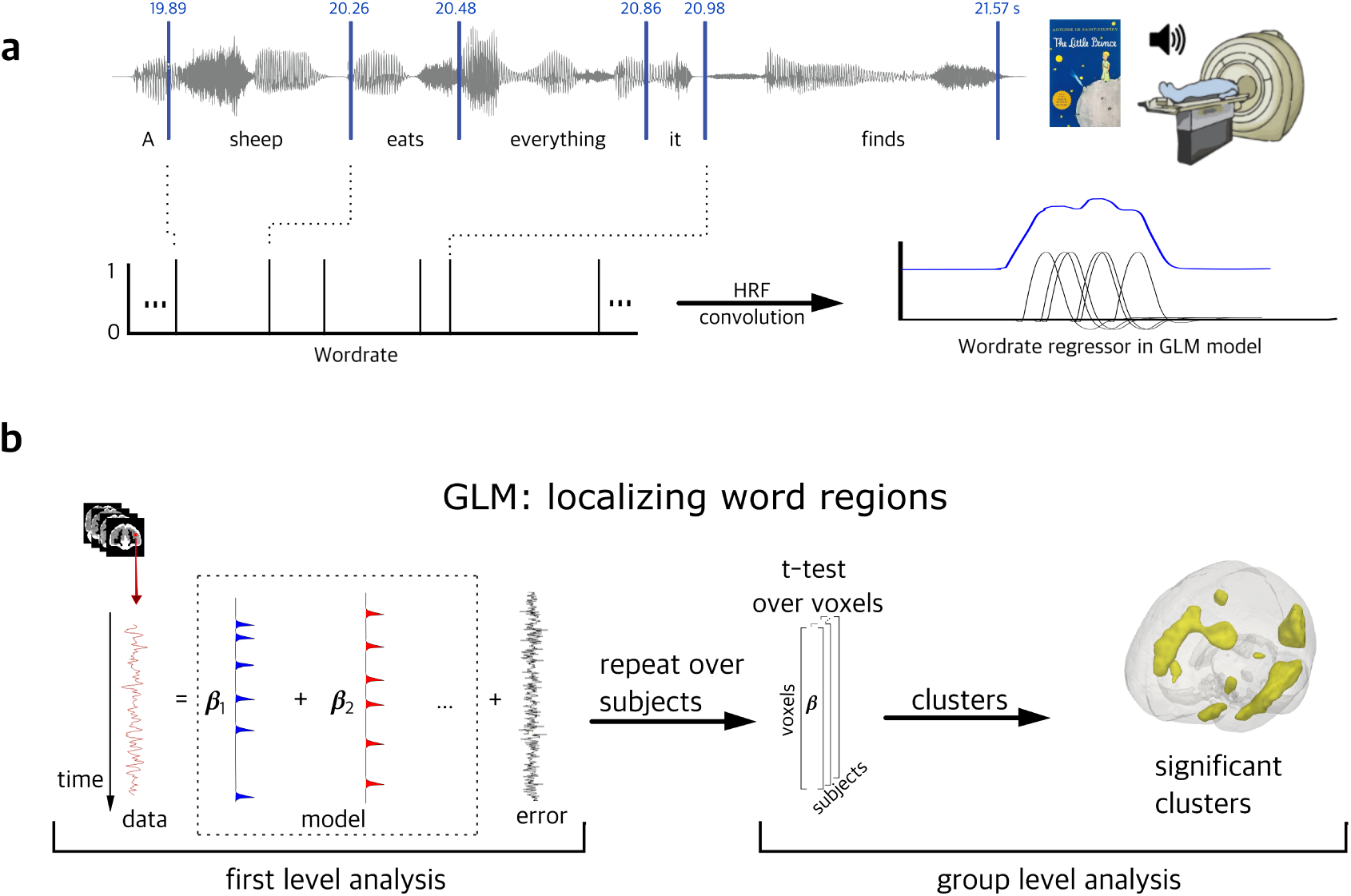
GLM analyses to localize the *wordrate* regressor. *a*. Offest of each word in the audiobook was marked 1 and was convolved with the canonical hemodynamic response function. *b*. The timecourse of each voxels BOLD signals was modeled using our designmatrix at the first level At the group level, a one-sample t-test was performed on the distribution of the beta values for the *wordrate* regressor across subjects at each voxel for the fMRI data. Statistical significance was held at *p <* .05 FWE with a cluster size greater than 50.

To illustrate the precise anatomical correspondence of our results with prior data, we overlaid fMRI term-based meta-analysis from Neurosynth^37^ (Retrieved September 2021) for the “pitch” area (https://neurosynth.org/analyses/terms/pitch/; from 102 studies) and the “words” area (https://neurosynth.org/analyses/terms/words/; from 944 studies). Our results are highly consistent with prior literature (see Figure 7). MNI coordinates of the significant clusters and their statistics are shown in Table 4.

**Figure 7.**
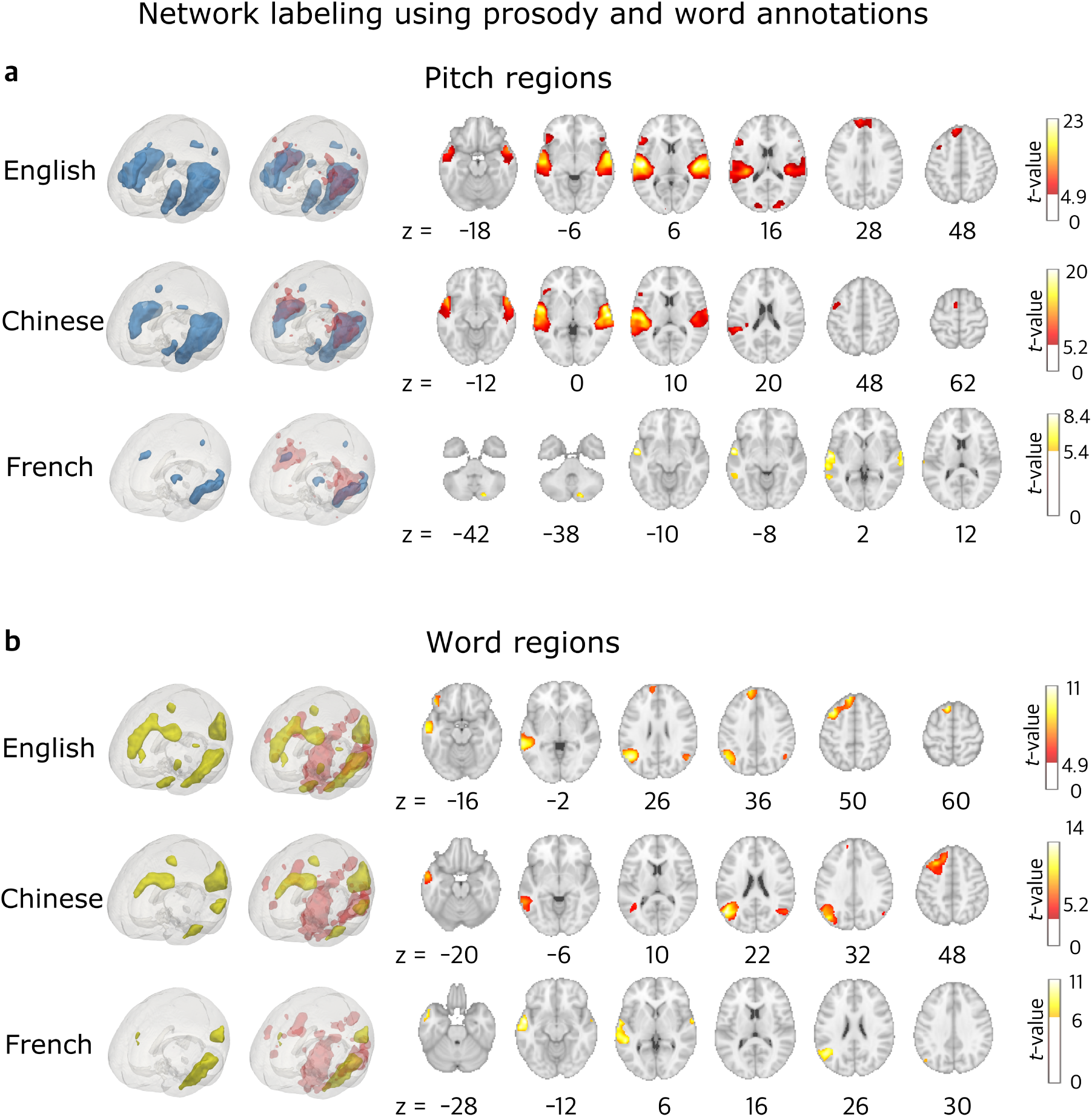
GLM results showing the significant clusters for **a**. the pitch and **b**. word regions in the English, Chinese and French data using *f0* and *wordrate* annotations. Red areas in the second column of the 3D brains shows meta-analyses of pitch and word regions from Neurosynth^37^. Statistical significance was thresholded at *p <* .05 FWE and *k >* 50.

**Table 4.** GLM results for the *f0* and *wordrate* regressors for the Chinese, English and French fMRI data: MNI coordinates, cluster size and their peak level statistics, thresholded at *p <* .05 FWE and *k >* 50.

| Condition | Language | Cluster | MNI Coordinates | k-size | t-value | p-value |
| --- | --- | --- | --- | --- | --- | --- |
| Prosody | Chinese | STG | 62,-14,0 | 3566 | 20.22 | <.001 |
|  |  | Heschl's Gyrus | -56,-6,4 | 5036 | 19.97 | <.001 |
|  |  | Frontal Lobe | -4,0,62 | 73 | 7.40 | <.001 |
|  |  | MFG | -52,-2,48 | 64 | 6.15 | 0.0005 |
|  | English | Heschl's Gyrus | -50,-18,6 | 5330 | 22.98 | <.001 |
|  |  | STG | 58,-20,4 | 5053 | 22.64 | <.001 |
|  |  | IFG | -52,26,10 | 864 | 10.09 | <.001 |
|  |  | SFG | -8,58,26 | 1272 | 9.55 | <.001 |
|  |  | Cerebellum | -18,-74,-32 | 245 | 8.84 | <.001 |
|  |  | Occipital Pole | 16,-94,16 | 188 | 7.95 | <.001 |
|  |  | Cerebellum | 22,-78,-36 | 245 | 7.40 | <.001 |
|  |  | MFG | -34,12,42 | 145 | 6.88 | <.001 |
|  |  | IFG | 52,26,-8 | 153 | 6.88 | <.001 |
|  |  | Occipital Pole | -14,-92,12 | 151 | 6.87 | <.001 |
|  | French | STG | -62,-12,4 | 1349 | 8.92 | <.001 |
|  |  | STG | 68,-22,2 | 218 | 7.09 | <.001 |
|  |  | Precuneus | -4,-70,32 | 53 | 6.63 | <.001 |
|  |  | MFG | -42,18,28 | 150 | 6.14 | <.001 |
| Word | Chinese | AG | -50,-64,22 | 2040 | 14.45 | <.001 |
|  |  | MFG | -28,22,48 | 1194 | 10.39 | <.001 |
|  |  | MTG | -56,0,-20 | 358 | 9.65 | <.001 |
|  |  | MTG | -60,-46,-6 | 511 | 8.61 | <.001 |
|  |  | AG | 54,-64,26 | 289 | 8.03 | <.001 |
|  | English | MTG | -52,-4,-28 | 1683 | 11.10 | <.001 |
|  |  | AG | -48,-60,26 | 1561 | 10.54 | <.001 |
|  |  | MFG | -36,16,50 | 1770 | 10.09 | <.001 |
|  |  | IFG | -46,32,-12 | 288 | 9.10 | <.001 |
|  |  | MTG | 60,-4,-30 | 171 | 6.92 | <.001 |
|  |  | IFG | -52,26,8 | 86 | 6.57 | <.001 |
|  |  | AG | 52,-64,28 | 191 | 6.55 | <.001 |
|  |  | Cerebellum | 18,-76,-30 | 82 | 5.52 | 0.010 |
|  | French | STG | -54,-4,-12 | 1674 | 9.45 | <.001 |
|  |  | AG | -50,-60,24 | 516 | 8.88 | <.001 |
|  |  | STG | 62,-2,2 | 72 | 7.01 | <.001 |

## Usage Notes

The LPPC-fMRI can advance our understanding of speech and language processing in the human brain during naturalistic listening. However, there are several limitations and usage bottlenecks, including annotations and analyses that we now discuss to help others use the LPPC-fMRI to make new discoveries.

### Annotation bottleneck

Most of the linguistic annotations were done automatically using existing NLP tools, which may contain errors and affect downstream annotations. For example, syntactic node counts for each word in the audiobooks based on bottom-up, top-down and left-corner parsing strategies were applied to the Stanford-derived constituency trees, and the accuracy of the tree structures will affect the number of node counts.

### Analysis bottleneck

Although GLM or encoding models have been commonly applied to fMRI data using long naturalistic stimuli like audio-books^8,9,11,18,38–40^. There are no standardised approaches for analysing complex and high dimensional naturalistic fMRI data. Machine learning approaches are becoming an increasingly common way to analyze fMRI data, and we encourage the development of innovative analysis approaches by running machine learning competitions on the LPPC-fMRI corpus.

## Code availability

Scripts used in this manuscript are available at https://github.com/jixing-li/lpp_data.

## Acknowledgements

This material is based upon work supported by the National Science Foundation under grant numbers 1903783 and 1607251, the French Agence Nationale pour la Recherche under grant NCM-NL ANR 16-NEUC-0005-02, and the Jeffrey Sean Lehman Fund for Scholarly Exchange with China at Cornell University. J.L. is supported by the NYU Abu Dhabi Institute under Grant G1001.

## Author contributions statement

J.H. designed the study. J.H., S.B. and J.L. collected and preprocessed the English data with the help of N.S. and W.L.. J.L. collected and preprocessed the Chinese data with the help of Y.Y.. C.P. collected and preprocessed the French data. W.L. provided data acquisition and preprocessing methods. J.L. and S.Z. did the technical validation of the data. B.F. prepared the OpenNeuro archive. J.L. wrote the manuscript with the help of J.H..

## Competing interests

The authors declare no competing interests.

## Notes

https://openneuro.org/datasets/ds003643/

### Competing Interest Statement

The authors have declared no competing interest.

